# A unified classification system for HIV-1 5’ long terminal repeats

**DOI:** 10.1101/2022.12.07.519241

**Authors:** Xing Guo, Dan Yu, Mengying Liu, Hanping Li, Mingyue Chen, Xinyu Wang, Xiuli Zhai, Bohan Zhang, Yanglan Wang, Caiqing Yang, Chunlei Wang, Yongjian Liu, Jingwan Han, Xiaolin Wang, Tianyi Li, Jingyun Li, Lei Jia, Lin Li

## Abstract

The HIV-1 provirus mainly consists of internal coding region flanked by the 2 same long terminal repeats (LTRs) at each terminus. The LTRs play important roles in HIV-1 reverse transcription, integration, and transcription by the association with host factors. However, despite of the significant study advances of the internal coding regions of HIV-1 by using definite reference classification, there are no systematic classifications for HIV-1 5’ LTRs, which hinders our elaboration on 5’ LTR and a better understanding of the viral origin, spread and therapy. Here, by analyzing all available resources of 5’ LTR sequences in public databases following 4 recognized principles for the reference classification, 83 representatives and 14 consensus sequences were identified as representatives of 2 groups, 6 subtypes, 6 sub-subtypes, and 9 CRFs. To test the reliability of our established classification system, the constructed references were applied to identify the 5’ LTR assignment of the 22 clinical isolates in China. The results revealed that 16 out of 22 tested strains showed a consistent subtype classification with the previous LTR-independent classification system. However, 6 strains, for which recombination events within 5’ LTR were demonstrated, unexpectedly showed a different subtype classification, leading a significant change of binding sites for important transcription factors including SP1, p53, and NF-κB. The binding change of these transcriptional factors would probably affect the transcriptional activity of 5’ LTR. This study established a unified classification system for HIV-1 5’ LTRs, which will facilitate HIV-1 characterization and be helpful for both basic and clinical research fields.

**IMPORTANCE:** Here, a scientific, reliable, and usable classification system based on the 5’ LTR sequences was established, which will allow us to effectively facilitate the precise typing of HIV-1 strains. This classification system was applied to 22 HIV-1 strains circulating in China, we found that 6 out of 22 strains analyzed, belonged to a different subtype when our results were compared to those obtained with the previous LTR-independent classification system. Thus, these data demonstrated that our classification method could greatly improve the HIV-1 subtype classification. We found that 6 5’ LTR sequences showed recombination events, leading to a significant exchange of the binding sites of transcriptional factors. Thus, this work established a comprehensive HIV-1 5’ LTR classification system, which will help the scientific community to precisely characterize HIV-1 variants, and better understand the origin and spread of HIV-1 strains, and it also may be helpful for pathogenicity and transmissibility evaluation studies.

## Introduction

Human immunodeficiency virus type 1 (HIV-1) is the most widespread type of HIV and has many subtypes, which have caused a complex problem for drug and vaccine design and development. HIV-1 provirus contains 2 same long terminal repeats (LTRs) at each terminus after integration. Each flanking viral LTR consists of U3 (455bp), R (96 bp), and U5 (83bp) (1, 2), contains elements of promoters, enhancers, TATA-independent promoters, and multiple transcription factor binding sites (TFBS) (3-5), and plays an important role in virus reverse transcription (6), integration (7-11), and transcription (5, 12, 13). Although both 5’ and 3’ LTRs have the function of Tat-induced promoter, the 5’ LTR element was the main transcriptional promoter, due to the fact that only the inactivation of a 5’ LTR promoter or lack of transcriptional function can trigger 3’ LTR transcription (12). A large number of transcription factors (TFs) have been shown to bind HIV-1 5’ LTR and regulate proviral transcription (5), such as nuclear factor-κB (NF-κB), nuclear factor for activated T cells (NFAT), specific protein 1 (SP1), TATA-binding protein (TBP), activator protein-1 (AP-1) (13, 14) and the positive transcriptional elongation factor b (P-TEFb) (15, 16).

Since the discovery of HIV-1, reference sequences for HIV-1 internal, 5’ LTR, or the full length has been proposed sporadically (2, 17-21). However, the unified reference guide to HIV-1 classification based on phylogenetics was systematically proposed by Robertson, D., et al., till 2000 by using the internal regions including *gag, pol, env*, and other 6 function regions (22). Taking advantage of these achievements, researchers have demonstrated that HIV-1 exhibit significant genetic diversity (6, 22, 23). Currently, HIV-1 has been divided into Main (M), Outlier (O), non-M/non-O (N), and P groups. M group is the most circulating group, which has been divided into 10 subtypes, 9 sub-subtypes, and 134 CRFs. In addition, the study in the classification of hepatitis C virus (HCV) has been also significantly improved since the systematic establishment of reference sequences by Simmonds et al. in 2005 and 2014 (24, 25).

The classification system mentioned above used the internal regions for HIV-1 classification without the involvement of 5’ LTRs sequences. Despite of the significant study advances of the internal regions of HIV-1 by using definite reference classification, there are no systematic classifications for HIV-1 5’ LTRs, which hinders our elaboration on 5’ LTR and a better understanding of the viral origin and spread (2, 17-19, 26). Intersubtype recombination in HIV-1 LTR has been demonstrated as an important viral evolutionary strategy, which also can affect the propagation of the virus (27, 28). However, current analysis tools such as the posterior probabilities-based HIV-1 recombination detection tool, jpHMM, are unable to provide accurate subtyping information of the 5’ LTR region (29, 30). Besides, the cis-acting elements and the various motifs of 5’ LTR have been widely studied for HIV-1 subtype B, but only a few works have focused on the other HIV-1 subtypes (18, 26, 31, 32) due to the unavailable of 5’ LTRs classification system.

With the increasing recognition of the important roles played by 5’ LTR, a systematic classification of 5’ LTR regions is needed. Therefore, we have proposed a unified classification system for HIV-1 5’ LTR and found that a considerable proportion of strains from patients samples showed distinct subtype assignment by using our system in comparison with the previous LTR-independent classification system, highlighting a critical role of LTR references for HIV-1 classification. This classification system will help to improve our research on HIV-1 epidemiology and evolution, and better understand the basic and clinical science fields of HIV-1.

## Materials and methods

### Sequences

All available 5’ LTR sequences of HIV-1 (any subtype) were retrieved from the HIV sequence database on August 17, 2021, including solo 5’ LTRs without the remaining complete genome and those 5’ LTRs with the remaining complete genome. The box of *one sequence/patient* was checked.

### Quality control

Multiple quality control strategies were performed to screen credible ones for reference construction and further analysis. Full-genome sequences assigned as unique recombinant form (URF) in the HIV sequence database were first excluded. The rest were submitted to the QC tool (https://www.hiv.lanl.gov/content/sequence/QC/index.html) and jpHMM (http://jphmm.gobics.de/) to validate the subtype assignment and perform further elimination. JpHMM tool is based on posterior probabilities and is rather intelligent. It can produce a genome subtype mosaic map. The recombination prediction in jpHMM is based on a precalculated multiple sequence alignment of the major HIV subtypes references. The evaluation of its prediction accuracy showed that it is more accurate than the competing methods used for recombination breakpoint detection (29, 30, 33). To ensure that the most accurate sequences can be screened for reference construction, those full-genome sequences with inconsistent subtype information were also excluded. Similarly, solo 5’ LTRs assigned as URF or without subtype information were excluded. The length of all the remaining sequences was further examined by comparison to reference HXB2 (accession number: K03455) to ensure high qualification. Finally, the screened high-quality sequences were preserved for subsequent comprehensive classification.

### The 5’ LTR determination of HIV-1 strains circulating in China

Here, plasma samples were collected from HIV-1-infected patients in Hebei province, China. These plasma samples were stored at -80 °C until use. Viral RNA was extracted using QIAamp^®^ Viral RNA Mini Kit (QIAGEN, 52904) according to the manufacturer’s instructions. Exactly following the natural reverse transcription process of HIV-1, the two partial LTRs at the 5’ terminus and 3’ terminus of genomic RNA were amplified, respectively, and then sequenced, assembled, and combined via the R region for a complete LTR (34). The primers in the NCR region and U3-R were designed. The nested PCR primers and the relative positions to HXB2 (accession number: K03455) were shown in Supplementary Table 1. The reverse transcription and the first round PCR were performed in a 25μl reaction volume by using PrimeScript™ One-Step RT-PCR Kit (TAKARA, RR055A). Cycling conditions were as follows: initial incubation at 50°C for 32 min, and at 94°C for another 3 min. Subsequently, 30 cycles were performed at 94°C for 30 sec, at 55°C for 30 sec, and at 72°C for 1 min followed by a final extension of 5 min at 72°C. We performed the second round of PCR using Premix Taq™ (TAKARA, RR902Q) in a 50μl reaction volume. Cycling conditions were as follows: initial incubation at 94°C for 3 min followed by 35 cycles at 94°C for 30 sec, at 60°C for 30 sec, and at 72°C for 1 min, a final extension at 72°C for 10 min. The PCR products were directly sequenced and sequences were assembled and edited by CONTING EXPRESS.

### Phylogenetic analysis

To confirm 5’ LTR references classification, the screened solo 5’ LTR sequences and 5’ LTRs with complete sequences were used to construct a maximum likelihood phylogenetic tree with MEGA X software, respectively (35). A maximum likelihood phylogenetic tree was also constructed to identify clinical isolates in Hebei province, China. The best-fitting models of nucleotide substitution were calculated by the model selection function in MEGA X. Tree topologies were searched using subtree-pruning-and-regrafting level 3 (SPR level 3), and the initial tree was made automatically (Default-NJ/BioNJ). The confidence of each node in phylogenetic trees was determined using the bootstrap test with 500 replicates. Bootstrap support values of ≥70% were considered to be significant. The final ML trees were visualized using iTOL v6 (36).

### Recombination analysis

There are 4 recognized requirements for subtype reference classification. The 4^th^ is the exclusion of intersubtypic recombination, whether the components were classified or not (22, 25). The reason is that compared to nucleotide substitution, recombination can dramatically change the content of genome and confuse the phylogenetic relationship (37, 38). To investigate whether the recombination events exist among the potential LTR references and the clinical sequences, RDP4 recombination analysis tool was used to perform systematic recombination analysis (39). RDP4 is a software package suitable for the statistical identification and characterization of recombination events in nucleotide sequences. It is very intelligent and can greatly improve both the sensitivity and reliability by using a series of nonparametric recombination detection methods: RDP, GENECONV, BOOTSCAN, MAXCHI, CHIMERA, SISCAN, 3SEQ, and LARD (40-47). Here, the highest acceptable *p*-value is set to 0.05. The sequence is set to linear. Other parameters are the default RDP4 settings. In order to ensure reliability, an HIV-1 5’ LTR sequence is considered to be recombinant when the recombinant signal is supported by at least four methods with the *p*-value≤ 0.05 after Bonferroni correction. The inferred breakpoint position is manually checked using the recombinant signal analysis implemented in RDP4. For details on the methods and algorithms of the recombination analysis tools used here, please refer to the comprehensive list of recombination analysis softwares maintained by Robertson laboratory (48).

### Specific TFBS identification in 5’ LTRs and TFs interaction predictions

The PROMO tool was used to find the characteristic TFBS in different subtypes and recombinants within 5’ LTRs. PROMO is a virtual laboratory for the identification of putative TFBS in DNA sequences from a species or groups of species of interest. TFBS defined in the TRANSFAC database are used to construct specific binding site weight matrices for TFBS prediction (49, 50).

### Ethics statement

The study involving human participants was reviewed and approved by the ethics committees of Beijing Institute of Microbial Epidemiology. All participants signed written informed consent prior to sample donations.

## Results

### Sequences

There are currently 129 identified groups, subtypes, sub-subtypes, and CRFs representatives in *Subtype Reference Alignments* provided in the HIV sequence database. They are widely recognized and used in the field, and therefore are named gold-standard sequences in the current work. Among them, A1, A4, A6, B, L, CRF02_AG, CRF03_A6B, CRF04_cpx, CRF06_cpx, CRF08_BC, CRF09_cpx, CRF11_cpx, CRF12_BF, CRF26_A5U, CRF27_cpx, CRF30_0206, CRF60_BC, O, and P subtypes contain the sequence of 5’ LTR. All of the remaining 110 representatives have no sequence of 5’ LTR, accounting for 85.27% (110/129) (Fig. 1a). Additionally, there are a total of 835,940 entries with classified subtypes by August 17, 2021. There are 803 5’ LTR entries by searching with any subtype, among which there are only 497 5’ LTR entries (one per patient) (Fig. 1b). There are 722 5’ LTR entries by searching with classified subtype term, among which there are only 431 5’ LTR entries (one per patient) (Fig. 1b). Then, a total of 497 5’ LTR sequences (any subtype) of HIV-1 were finally retrieved from HIV sequence database (Supplementary Dataset 1), which were divided into two subsets. One subset included 411 5’ LTR sequences, which contained the remaining genome region of HIV-1. Another subset included 86 5’ LTR sequences with no region of the remaining HIV-1 genome, which were defined as solo 5’ LTR.

**Figure 1.**
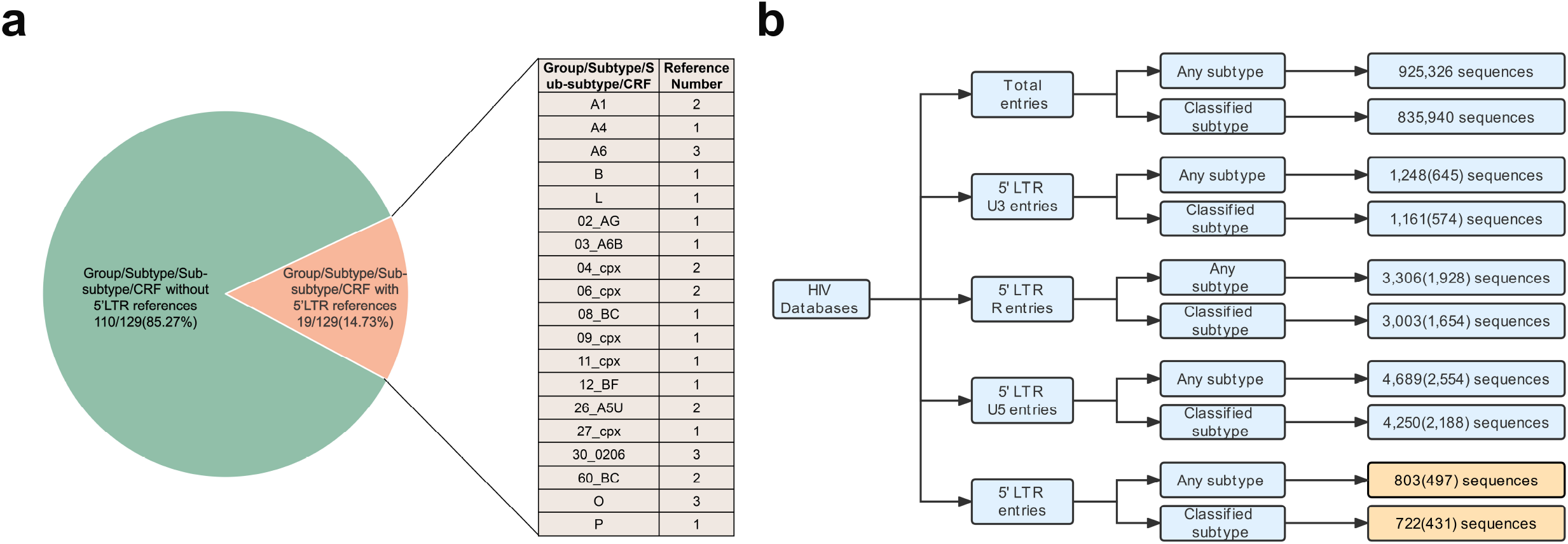
Inclusion of 5’ LTR region in HIV sequence database. (a) Completeness of 5’ LTR region in recognized *Subtype Reference Alignments* provided in the HIV sequence database. By August 17, 2021, among 129 the currently identified groups, subtypes, sub-subtypes, and circulating recombinant forms (CRFs) representatives, only A1, A4, A6, B, L, CRF02_AG, CRF03_A6B, CRF04_cpx, CRF06_cpx, CRF08_BC, CRF09_cpx, CRF11_cpx, CRF12_BF, CRF26_A5U, CRF27_cpx, CRF30_0206, CRF60_BC, O, and P have 5’ LTR region. (b) Entries summary of HIV-1 sequences in the Los Alamos National Laboratory HIV sequence database by August 17, 2021.

### Quality control results

To improve the reliability of the reference classification, multiple quality control strategies have been performed. Defective sequences with low quality were excluded. As for full-genome sequences, 44 out of 411 were first excluded for being assigned as URFs in the HIV sequence database. The remaining 367 full-length sequences were submitted to the Quality Control tool and jpHMM to further validate assignment accuracy. A total of 36 sequences were removed for no feedback by jpHMM. A total of 26 sequences were eliminated for subtype inconsistency. 3 sequences belonging to uncertain subtype definitions were also excluded. 27 sequences belonging to group O and 1 sequence belonging to group P were preserved as outgroup. As for solo 5’ LTRs, 5 sequences assigned as URFs in the HIV sequence database were excluded and another 2 solo 5’ LTRs were excluded for no subtype information. 1 sequence belonging to U was excluded. Subsequently, the length of all sequences was further examined by comparison to reference HXB2 (accession number: K03455). Another 2 solo 5’ LTRs were excluded due to excessive length (2185 bp and 1920 bp) resulting from insertions. The details of excluded sequences are summarized in Supplementary Table 2. Finally, a total of 302 5’ LTRs with full-length sequences and 76 solo 5’ LTR sequences without complete genome were screened for further study (Supplementary Dataset 2 and Supplementary Dataset 3).

### Provisional 5’ LTR reference classification from gold-standard sequences

There are 4 recognized requirements for subtype references classification: (1) one or more complete sequence(s); (2) at least three epidemiologically unrelated isolates; (3) distinct clusters in a phylogenetic tree; (4) exclusion of intersubtypic recombination, whether the components were classified or not (22, 24, 25). Because sequences included in *Subtype Reference Alignments* have been strictly screened and recognized as standard references and widely used in related research of the field (51-53), they were named gold-standard sequences in this study. We found that among the 302 screened 5’ LTRs with full-length sequences, 20 sequences belong to gold-standard sequences (Supplementary Dataset 4). 5’ LTRs of these 20 gold-standard sequences have great potential to become high-quality reference sequences of the corresponding subtype and thus were provisionally assigned as the references for each subtype (Supplementary Table 3).

### Reference classification based on solo 5’ LTRs sequences

The screened 76 solo 5’ LTRs were used to construct a maximum likelihood phylogenetic tree by using MEGA X to identify distinct clusters, together with all the 20 provisional 5’ LTR references identified from gold-standard sequences. The best-fitting model of nucleotide substitution was generated and calculated as GTR+G+I by using MEGA X. The sequence (accession number: AF196750) was first eliminated for subsequent classification because it is assigned as subtype A1 while according to the phylogenetic tree it is displayed as CRF02_ AG in the HIV sequence database. Besides, the phylogenetic tree clearly showed confirmed distinct subtype clusters of all sequences including provisional 5’ LTR references from gold-standard sequences (Fig. 2).

**Figure 2.**
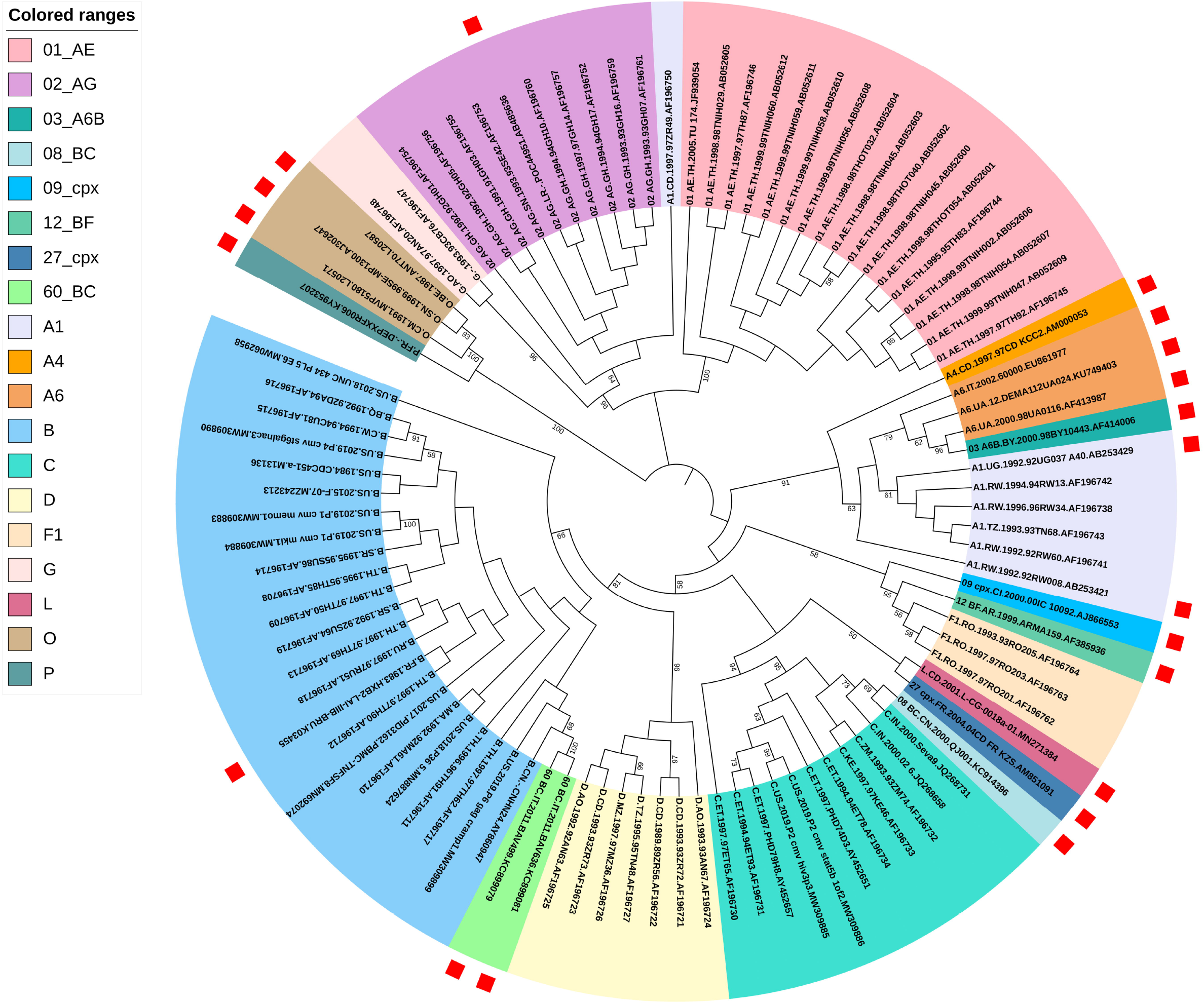
The ML phylogenetic tree built using the screened 76 solo 5’ LTRs sequences. The phylogenetic tree clearly showed confirmed distinct subtypes clusters, all supported by very strong bootstrap values. The best-fitting model of nucleotide substitution was GTR+G+I by using MEGA X. Tree topologies were searched using subtree-pruning-and-regrafting level 3 (SPR level 3) and the initial tree was made automatically (Default-NJ/BioNJ). The confidence of each node in the phylogenetic tree was determined using the bootstrap test with 500 replicates and values below 50% are not shown. The red squares represent 20 references identified from gold-standard sequences.

Considering countries and time distribution as well as the position relative to gold-standard sequences, 22 solo 5’ LTRs were assigned as the corresponding subtype references separately (Supplementary Table 3, Supplementary Dataset 5), which were classified into 4 subtypes, 2 sub-subtypes, and 2 CRFs (Supplementary Table 3).

### Reference classification based on 5’ LTRs with complete genomes

A total of 302 5’ LTRs sequences with complete genome were obtained after screening. Here, these full-length sequences together with all 20 provisional 5’ LTRs references carrying complete genome from gold-standard sequences were used to perform a maximum likelihood phylogenetic analysis using MEGA X. The best-fitting model of nucleotide substitution was generated and calculated as GTR+G+I by MEGA X. The results showed that sequences clustered in subtypes-specific branches with very high bootstrap values (Fig. 3a). Subsequently, 5’ LTRs sequences were extracted from the complete genome sequences to perform another round of maximum likelihood phylogenetic analysis to verify the clustering (Fig. 3b). The best-fitting model of nucleotide substitution was generated and calculated as GTR+G+I using MEGA X. The two phylogenetic trees were compared and sequences with inconsistent clustering were excluded. As shown in Fig. 3a and b, the sequences belonging to subtype B, C, D, G, H, F1, CRF07_BC, CRF08_BC, CRF12_BF, group O, and P consistently clustered in both trees. Thus, these sequences are all preserved.

**Figure 3.**
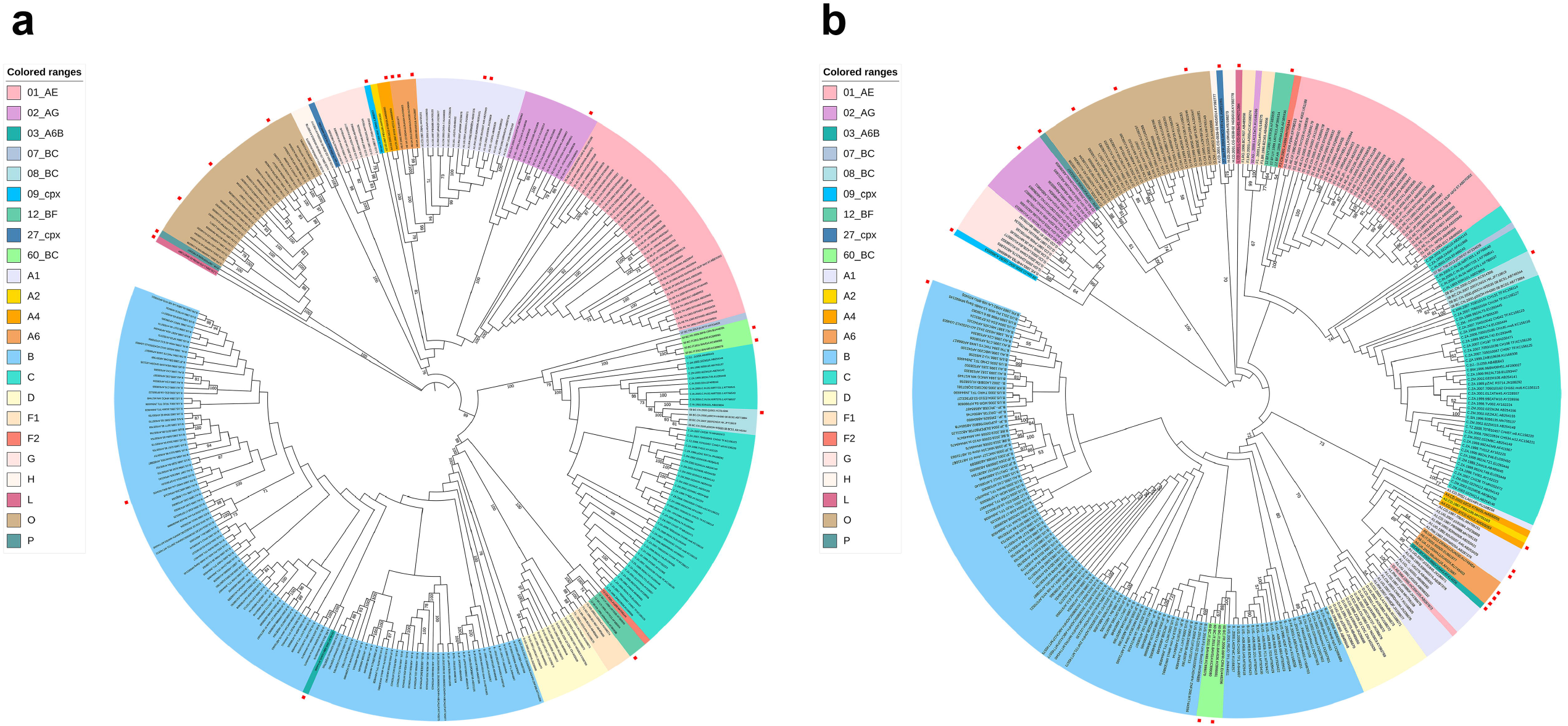
The ML phylogenetic analysis using 302 full-length genomes and 5’ LTRs extracted from them, respectively. (a) The maximum likelihood phylogenetic analysis by MEGA X based on 302 full-length sequences containing 5’ LTRs sequences. Sequences clustered in subtypes-specific branches with very high bootstrap values. (b) The maximum likelihood phylogenetic analysis by MEGA X based on extracted 5’ LTRs derived from the 302 full-length sequences. The confidence of each node in phylogenetic trees was determined using the bootstrap test with 500 replicates and values below 50% are not shown. Colored ranges represent different groups, subtypes, sub-subtypes, and CRFs. The red squares represent 20 provisional references identified from gold-standard sequences.

All 4 sequences of CRF60_BC, including 2 from gold-standard representatives (accession number: KC899079 and KC899081), formed a distinct cluster on each tree, with strong bootstrap values (Fig. 3a,b). However, the two clusters displayed in inconsistent topology. They nested within the subtype C clusters at the full-length analysis but nested within subtype B at the 5’ LTR level. We found that the reason is due to the fact that these 5’ LTRs sequences have been characterized to contain a large fragment of subtype B (27). Thus, these four sequences have also been preserved. Similarly, CRF03_A6B (accession number: AF414006) also formed a distinct cluster on each tree but displayed an inconsistent topology (Fig. 3a,b). It nested within the subtype B clusters at the full-genome analysis but nested within subtype A6 at the 5’ LTR level. The 5’ LTR has been previously characterized to be A6 (54). Thus, this CRF03_A6B has also been preserved. In addition, according to the definition of the recombinant structure by Vidal et al. (28), the CRF27_cpx (accession number: AM851091, the gold-standard sequence) with inconsistent topology between two trees was thus preserved.

Moreover, the phylogenetic analysis showed that sub-subtype A1, A2, A4, and A6 displayed a certain degree of topology dissociation. Considering the higher intra-subtypes sequence homology than the inter-subtypes sequence homology and that the dissociation occurred just within the whole cluster A, they were preserved as well.

It should be noted that there is the discrepancy of subtype L (accession number: MN271384, gold-standard sequence), sub-subtype F2 (accession number: MH705144), and CRF09_cpx (accession number: AJ866553, gold-standard sequence) and since no more than one sequence for each was available, they were all preserved but they were indicated with the asterisk (*), indicating risk warning as shown in Table 1.

**Table 1.**
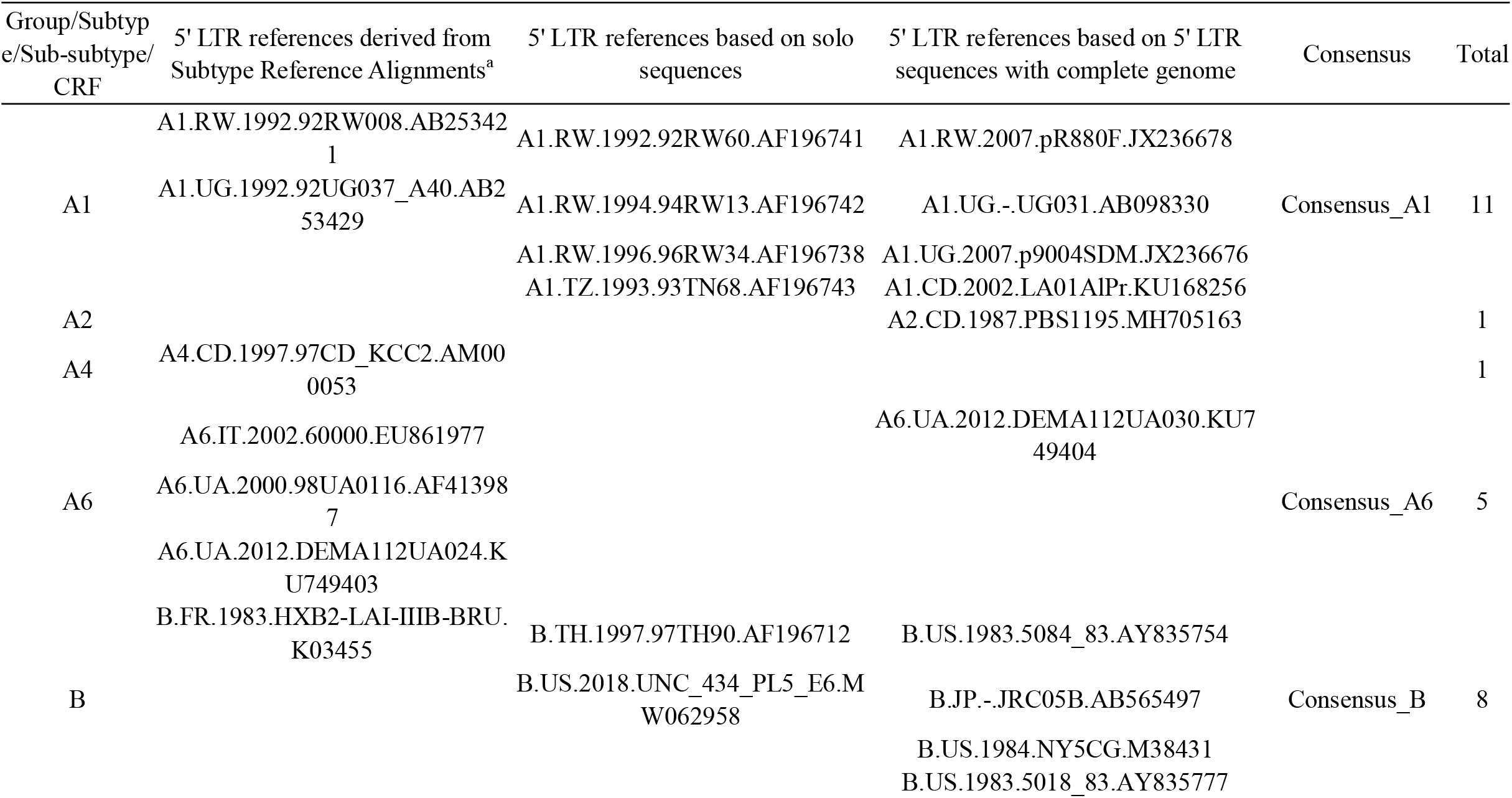
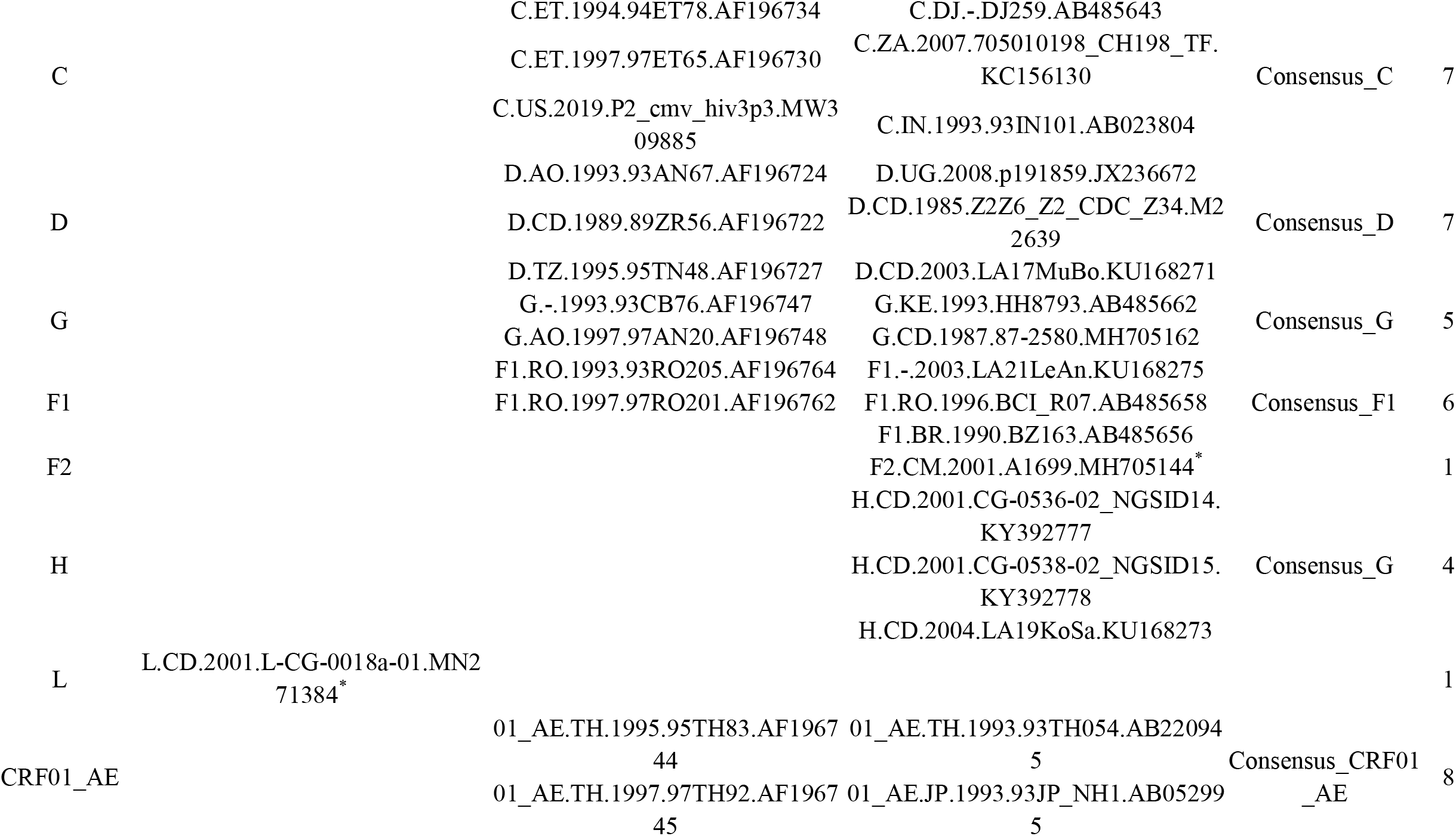
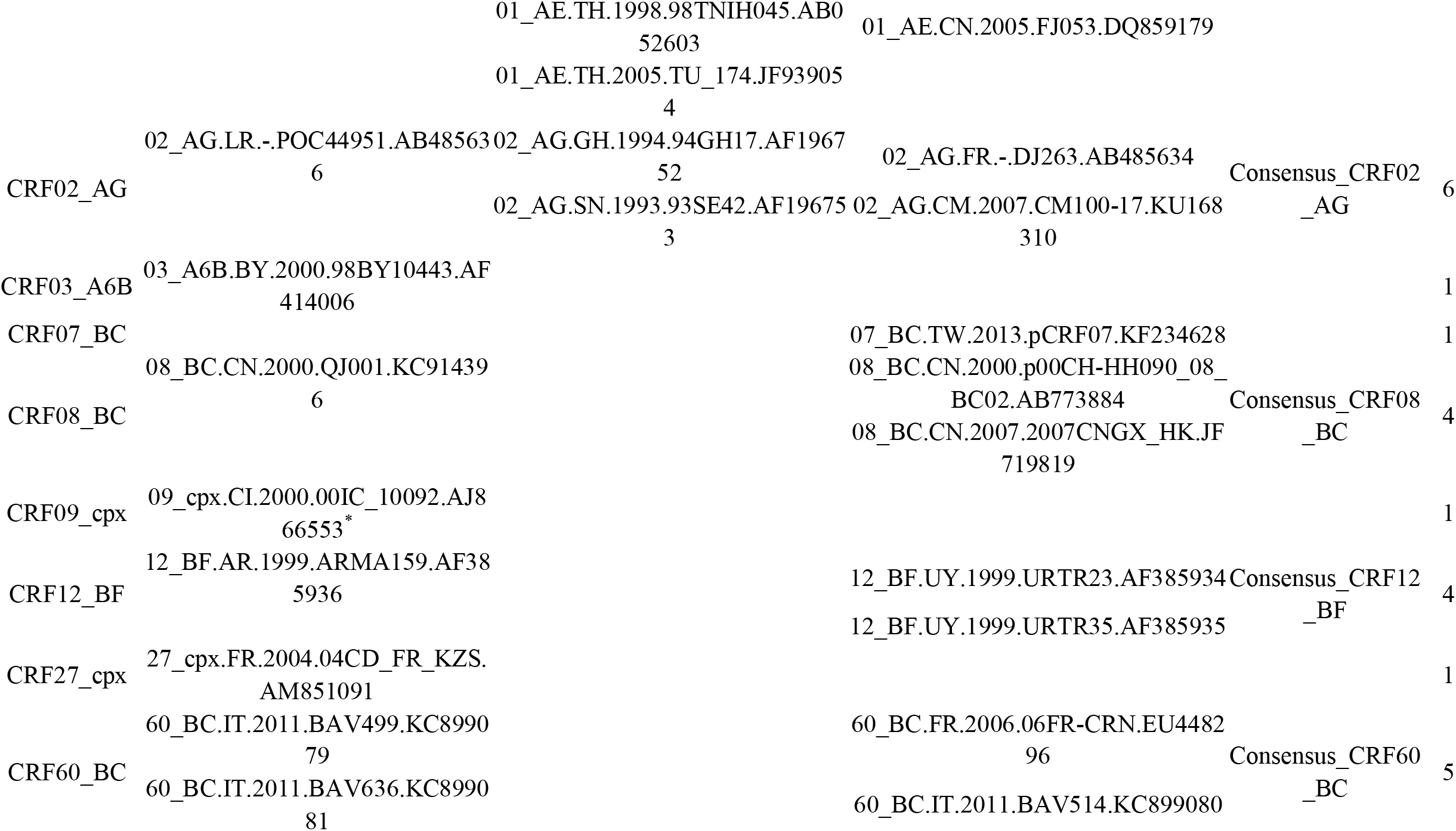
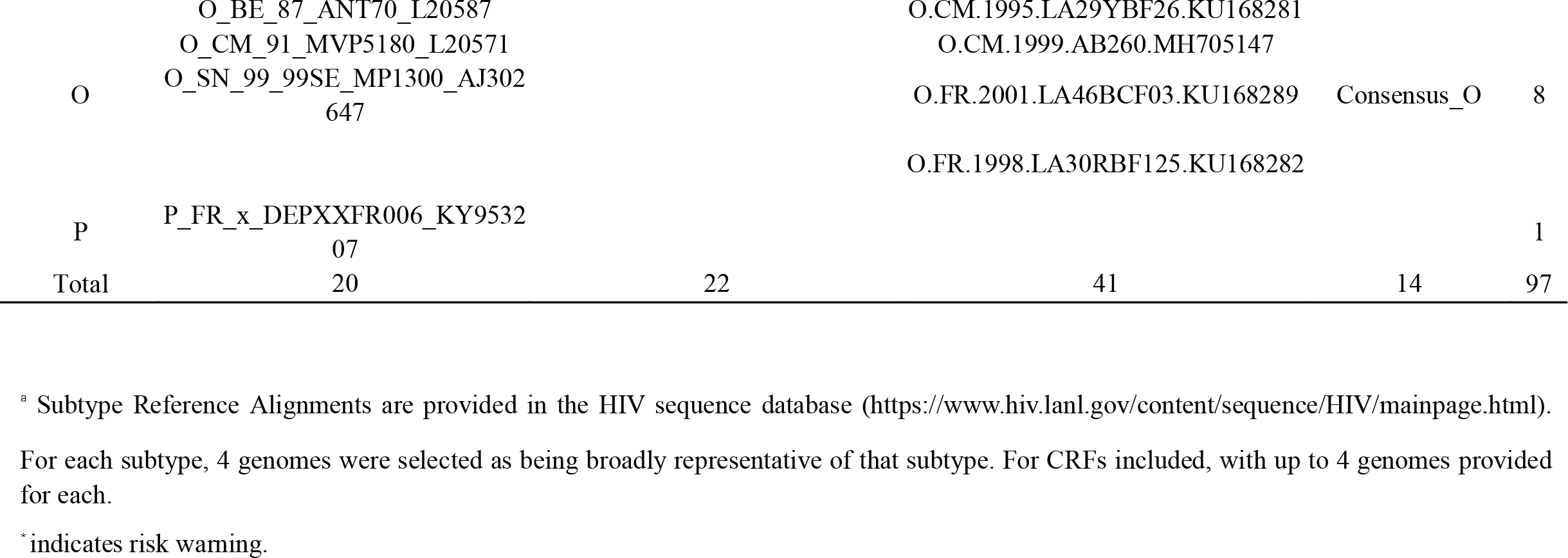
5’ LTR reference sequence statistics

From the comparison of the two phylogenetic trees conducted in this study, we found that one sequence (accession number: AB097872) nested within CRF01_ AE cluster at the full-genome level, but the 5’ LTR clustered with A1, in contrast with other CRF01_AE sequences. The remaining sequences all consistently clustered in the same manner in both trees. Thus, this sequence (accession number: AB097872) was excluded. Besides, one sequence belonging to CRF02_AG (accession number: KU168266) clustered within CRF02_AG in the complete genome-based analysis, as expected. However, its 5’ LTR region clustered to F1 subtype. This sequence was also removed.

Considering countries and time distribution as well as the position relative to gold-standard sequences, a total of 43 5’ LTR sequences with complete genome were assigned as the corresponding subtype references separately (Supplementary Table 3), which were classified into 1 group (O), 5 subtypes, 6 sub-subtypes, and 6 CRFs (Supplementary Table 3).

### Definitive determination of reference strains and genetic analysis of HIV-1 5’ LTRs sequences

One of the 4 recognized principles for the HIV-1 subtype classification is the absence of intersubtype recombination (22, 24, 25). Thus, the tool of detecting recombination, RDP4 (55), was applied to investigate whether any recombination occurred among the 85 5’ LTR references sequences. It was found that a sequence of subtype G (accession number: MH705145) and a sequence belonging to subtype A4 (accession number: AM000055) are recombinants. The sequence of subtype G (accession number: MH705145) is a recombinant result from unknown (major parent) and A1 (accession number: AF196741, minor parent) with the recombination signal supported by 7 recombination methods (RDP, GENECONV, BootScan, Maxchi, Chimaera, SiSscan, and 3Seq). The sequence of sub-subtype A4 (accession number: AM000055) was a recombinant from A1 (major parent, accession number: AB098330) and an unknown sequence (minor parent) with the recombination signal supported by 5 recombination methods used (RDP, BootScan, MaxChi, SiSscan, and 3Seq). Based on the 4^th^ principle of the reference system construction (22, 25), these two sequences were removed from the classified reference sequences evaluated in this study.

Therefore, a total of 83 subtype references covering 2 groups, 6 subtypes, 6 sub-subtypes, and 9 CRFs were selected as reference representatives (Table 1 and Supplementary Dataset 6). Then, these representative sequences were aligned and used for constructing a phylogenetic tree. Despite the length of the 5’ LTR region is short (634 bp) and the nucleotide identity is much higher than that of HIV-1 internal region, distinct clusters were still evident, even within sub-subtypes (Fig. 4). There is only one representative reference for subtype A2, A4, F2, L, CRF03_A6B, CRF07_BC, CRF09_cpx, CRF27_cpx, and group P. For subtype H, there formed a separate cluster but with low bootstrap support. On the contrary, all other clusters showed high bootstrap support. Further, 14 consensus sequences of 5’ LTR references were also constructed (Table 1, Supplementary Dataset 7).

**Figure 4.**
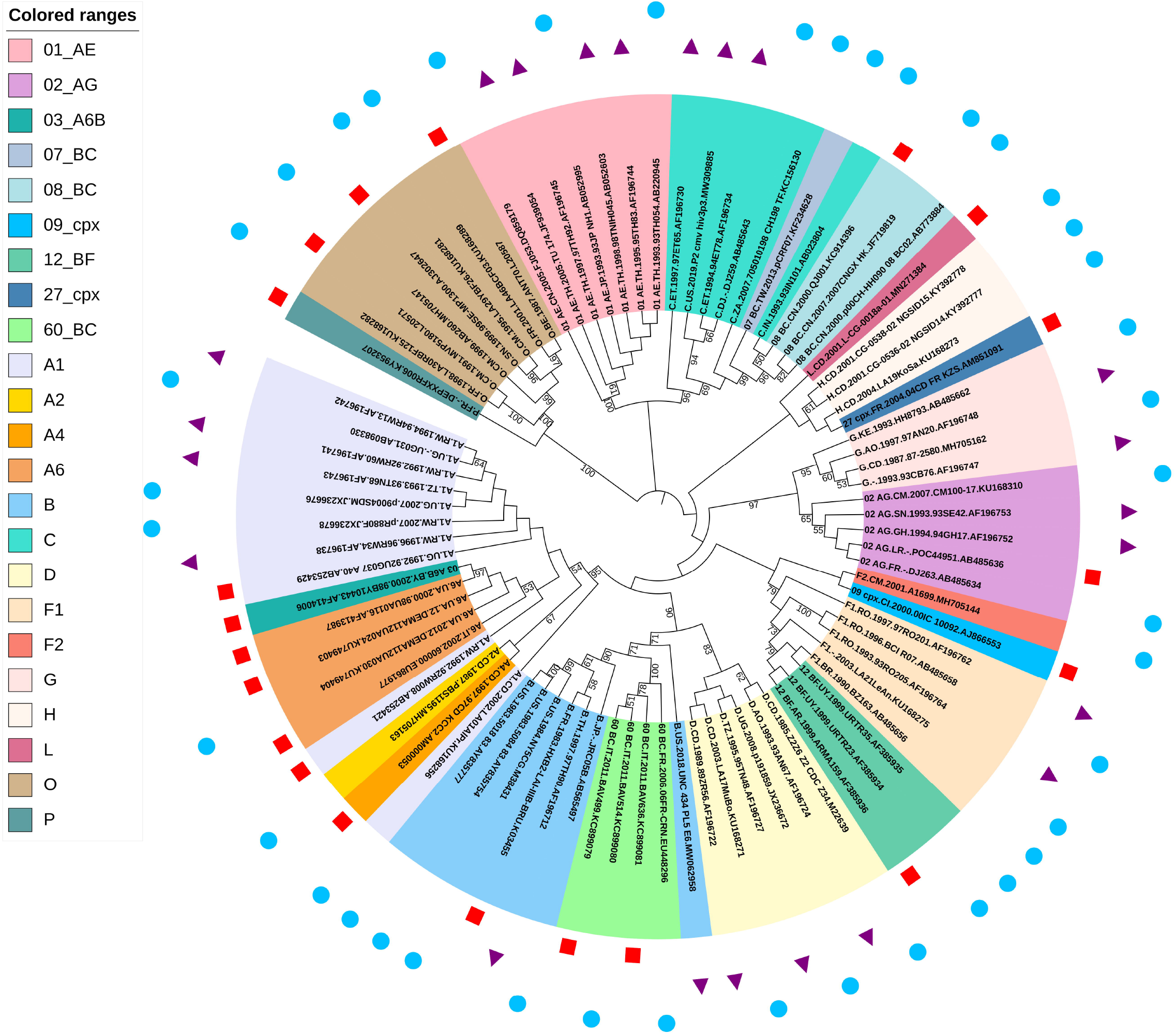
The phylogenetic tree of 83 definitive screened reference representatives of HIV-1 5’ LTRs. Representatives of each confirmed group, subtype, sub-subtype, and CRF were aligned and a phylogenetic tree was constructed using the maximum likelihood method by MEGA X. The results indicated that major distinct clusters are all with reliable bootstrap support and values below 50% are not shown (replicates=500). Tips are labeled by groups/subtypes/sub-subtypes/CRFs, sample country, sample year, sequence name, and accession number. Colored ranges represent different groups, subtypes, sub-subtypes, and CRFs. The red squares represent 20 references identified from gold-standard sequences. The purple triangles represent 22 references identified from solo 5’ LTRs. The blue dots represent 41 references identified from 5’ LTR sequences with complete genome.

It is worth to be noted that recombination exclusion analysis showed that the 5’ LTRs sequences belonging to 5 CRF02_AG representatives are all recombinants with the same mosaic manner (Fig. 5a). Specifically, the 5’ LTR sequence of sub-subtype A1 was inserted into the skeleton of the 5’ LTR sequence of subtype G at the position of 278-423bp (position according to the K03455 coordinates) (Fig. 5a). To further confirm the recombination analysis results, the fragments of parental strains were extracted and sub-region phylogenetic trees were constructed with parental strains using MEGA X. The results showed that the major parental strains belonged to subtype G with bootstrap support values of 91% (Fig. 5b), while the minor parental strains clustered within sub-subtype A1 (Fig. 5b). The length of the segment was only 146 bases, and as discussed by Leitner et al. (56), region length is not sufficient to produce phylogenetic trees with high bootstrap values, and similarly to previously reported, a high bootstrap value was not obtained in this study either, despite a distinct cluster still obtained. Therefore, it was clearly demonstrated in this study that the 5’ LTR sequences belonging to CRF02_AG subtypes were all recombinants.

**Figure 5.**
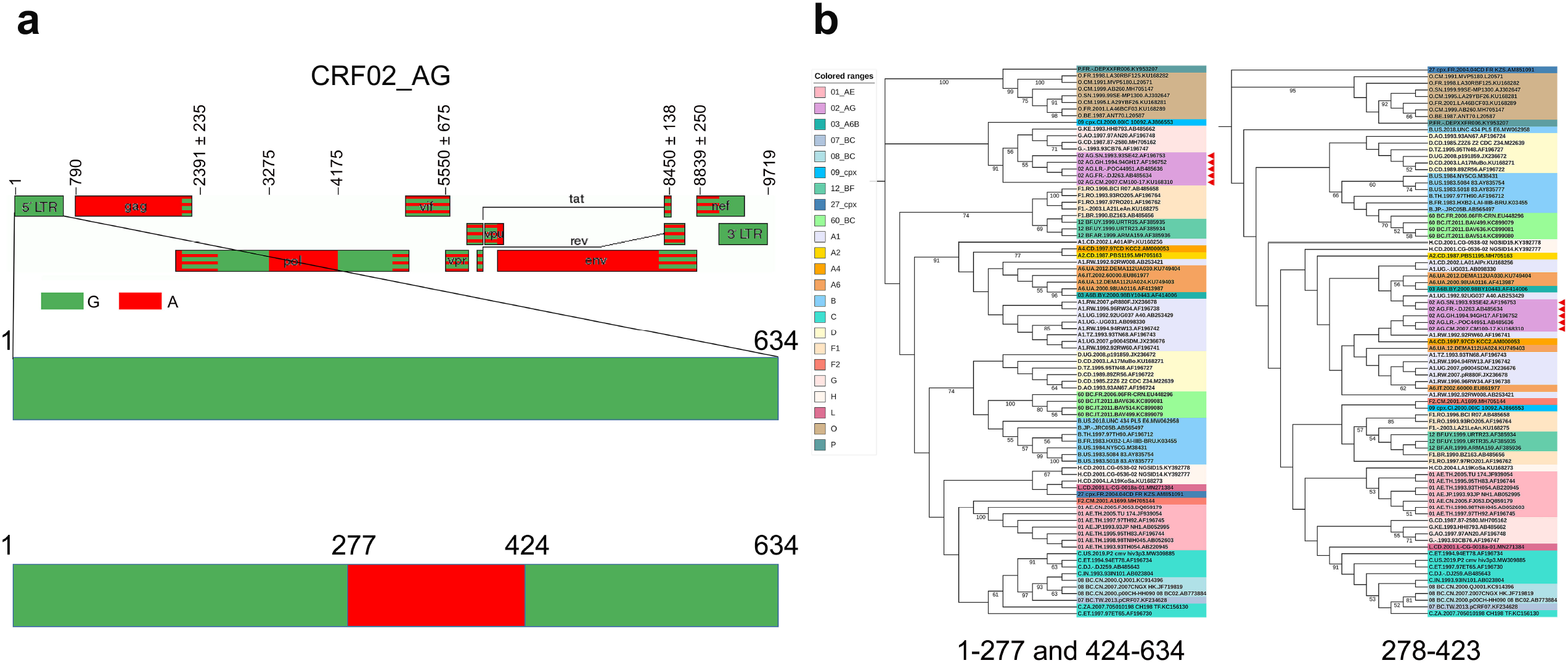
The recombination characterization of 5’ LTR of CRF02_AG references. The recombination analysis and phylogenetic analysis confirm that the 5’ LTRs of all 5 CRF02_AG representatives are recombinants with the same mosaic manner (including 02_AG.LR.-.POC44951.AB485636 from gold standard representatives in *Subtype Reference Alignments*). (a) Updated genome maps of CRF02_AG 5’ LTR. The top picture shows the original mosaic structure from the Los Alamos HIV sequence database. The below picture displays the updated recombinant details of 5’ LTR. The standard representatives are marked by different clors, as indicated. (b) The phylogenetic confirmation of the recombination pattern by sub-region trees using extracted fragments. The left picture shows the phylogenetic relationship of the region spanning HXB2 nt 1-277 and 424-634bp with all classified 5’ LTRs representatives in the current work. The right picture shows the phylogenetic relationship of the region spanning HXB2 nt 278-423bp with all classified 5’ LTRs representatives in the current work. The tree was constructed using the Maximum likelihood method implemented in MEGA X. The values at the nodes indicate the percent bootstraps in which the cluster to the right was supported. Colored ranges represent different groups, subtypes, sub-subtypes, and CRFs. The red triangles represent recombinants.

### Reliability test of newly established 5’ LTR classification system

To test the reliability and applicability of our established classification system, the constructed references were first applied to identify the 5’ LTR assignment of the clinical isolates in China. Genes of *gag, pol*, and *env* per isolate were amplified by RT-PCR, sequenced, and assembled as described in previous works (57-59). In addition, the two partial LTRs sequences at the 5’ terminus and 3’ terminus were also amplified, sequenced, assembled, and further combined via the R region to obtain a complete LTR sequence, as previously described (6). Thus, a total of 22 sequences including 4 regions (LTR, *gag, pol*, and *env*) were obtained.

Subsequently, a subtyping test of these 22 sequences by using the 3 regions (*gag, pol*, and *env*) was performed as previously described (57-59), followed by a subtyping test based on the 5’ LTRs which was performed using the classification system developed in this study. The results showed that 5’ LTRs of 16 out of 22 displayed consistent subtype assignment with the subtype assignment by using 3 other regions (*gag, pol*, and *env*). These subtypes included 2 subtype B, 8 CRF01_AE, 1 CRF07_BC, 2 CRF08_BC, and 3 CRF55_01B (Table 2, Fig. 6a). Unexpectedly, 3 sequences showed different subtype assignments in the 5’ LTRs compared to that obtained with the other 3 regions. HB020068 was demonstrated to belong to subtype CRF07_BC based on *gag, pol*, and *env* regions, but it was classified as subtype B based on the 5’ LTR region. On the contrary, HB070040 and HB070063 belonged to subtype B based on *gag, pol*, and *env* regions, but to CRF07_BC based on 5’ LTR regions (Table 2, Fig. 6a). The results revealed that the whole 5’ LTR of all 3 sequences was exchanged via genetic recombination. Furthermore, the other 3 sequences showed an exchange of a certain proportion of fragments within the 5’ LTR via recombination, including HB010151, HB010161, and HB030133. The recombinant HB010151 was produced by the recombination between HB010154 (CRF07_BC) and HB010165 (CRF01_AE). A segment of 455 bp of CRF07_ BC was inserted at positions 1-455 (position according to the K03455 coordinates) in the skeleton of CRF01_AE. HB010161 sequence displayed the same recombination events as HB010151 (Fig. 6b). The recombinantHB030133 resulted from the recombination between HB070068 (subtype B) and HB030138 (CRF08_BC). A segment of 207 bp of CRF08_ BC was inserted at positions 428-634 (position according to the K03455 coordinates) within the skeleton of subtype B (Fig. 6c). The mosaic structures were all confirmed by sub-region phylogenetic trees by using the fragments extraction (Fig. 6b,c). These results validated the accuracy of the classification system and its practical applicability and demonstrated that the whole classification system could facilitate the characterization of HIV-1 5’ LTRs.

**Table 2.** Assignments of 4 regions of the 22 amplified samples

| Sample | Assignment of *gag* | Assignment of *pol* | Assignment of *env* | Assignment of LTR |
| --- | --- | --- | --- | --- |
| HB070049 | B | B | B | B |
| HB070068 | B | B | B | B |
| HB010155 | CRF01_AE | CRF01_AE | CRF01_AE | CRF01_AE |
| HB010165 | CRF01_AE | CRF01_AE | CRF01_AE | CRF01_AE |
| HB010181 | CRF01_AE | CRF01_AE | CRF01_AE | CRF01_AE |
| HB010183 | CRF01_AE | CRF01_AE | CRF01_AE | CRF01_AE |
| HB010189 | CRF01_AE | CRF01_AE | CRF01_AE | CRF01_AE |
| HB010192 | CRF01_AE | CRF01_AE | CRF01_AE | CRF01_AE |
| HB020076 | CRF01_AE | CRF01_AE | CRF01_AE | CRF01_AE |
| HB020078 | CRF01_AE | CRF01_AE | CRF01_AE | CRF01_AE |
| HB010154 | CRF07_BC | CRF07_BC | CRF07_BC | CRF07_BC |
| HB030138 | CRF08_BC | CRF08_BC | CRF08_BC | CRF08_BC |
| HB040020 | CRF08_BC | CRF08_BC | CRF08_BC | CRF08_BC |
| HB030215 | CRF55_01B | CRF55_01B | CRF55_01B | CRF55_01B |
| HB070073 | CRF55_01B | CRF55_01B | CRF55_01B | CRF55_01B |
| HB010053 | CRF55_01B | CRF55_01B | CRF55_01B | CRF55_01B |
| HB020068 | CRF07_BC | CRF07_BC | CRF07_BC | B |
| HB070040 | B | B | B | CRF07_BC |
| HB070063 | B | B | B | CRF07_BC |
| HB010151 | CRF01_AE | CRF01_AE | CRF01_AE | CRF07_BC&CRF01_AE |
| HB010161 | CRF01_AE | CRF01_AE | CRF01_AE | CRF07_BC&CRF01_AE |
| HB030133 | B | B | B | B&CRF08_BC |

**Figure 6.**
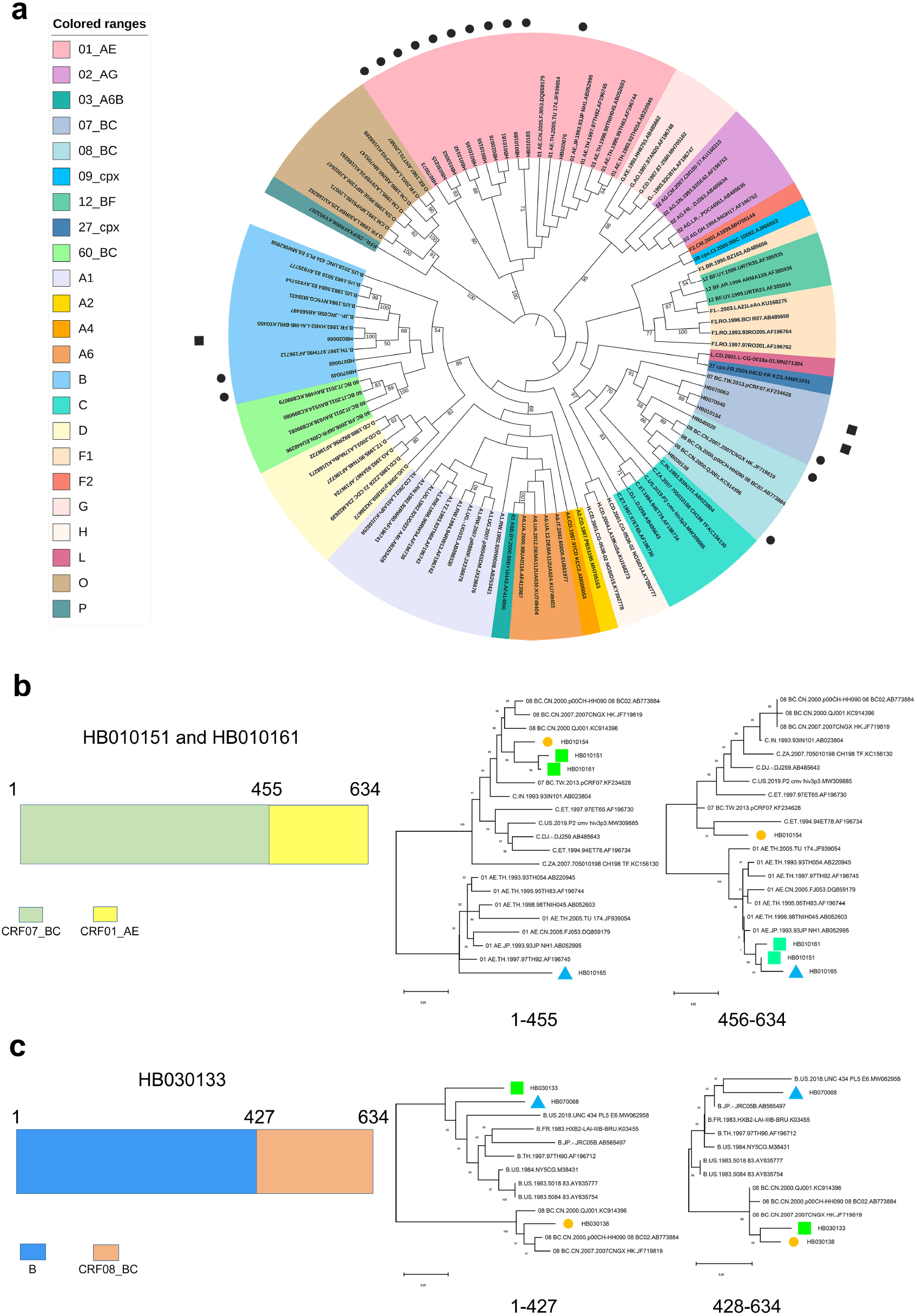
The reliability test of newly established 5’ LTR classification system by identifying the 5’ LTR assignment of the clinical isolates in China. (a) The ML phylogenetic tree was built using 19 amplified HIV-1 5’ LTR sequences together with 83 HIV-1 5’ LTR references. These 19 5’ LTR contained no recombination and showed good clustering with strong bootstrap support. The black dot represents 16 5’ LTRs having consistent subtype assignment with that of the 3 other regions including *gag, pol*, and *env*. The black square represents 3 5’ LTRs having inconsistent subtype assignments with that of the 3 other regions. Colored ranges represent different groups, subtypes, sub-subtypes, and CRFs. (b) The same recombination pattern of the 2 5’ LTR recombinants (HB010151 and HB010161) and the subsequent confirmation by sub-region trees. The 2 recombinants share a common mosaic pattern. The sub-region phylogenetic analysis confirmed that the extracted segments have the right subtype assignment as indicated by recombination analysis. The green square represents recombinants. The blue triangle represents major parent. The yellow dot represents minor parent. (c) The recombination pattern of HB030133 and the subsequent confirmation by sub-region trees. The sub-region phylogenetic analysis confirmed that the extracted segments have the right subtype assignment as indicated by recombination analysis. The green square represents recombinant. The blue triangle represents major parent. The yellow dot represents minor parent.

### Recombination in 5’ LTRs caused a significant change of transcription factor binding sites

We further explored the TFBS differences within 5’ LTRs of 6 identified recombinants circulating in China, including HB020068, HB070040, HB070063, HB010151, HB010161, and HB030133. The results revealed that recombination introduces extensive changes of a large number of TFBSs. For the exchange of 5’ LTR between subtype B and CRF07_BC (HB020068, HB070040, and HB070063), the exchange via recombination leads to a change of 1011 TFBSs (Supplementary Table 4). For the exchange of partial 5’ LTR between CRF01_AE and CRF07_BC (HB010151 and HB010161, Fig. 6b), the exchange via recombination leads to a change of 1326 TFBSs (Supplementary Table 5). For the exchange of partial 5’ LTR between subtype B and CRF08_BC (HB030133, Fig. 6c), the exchange via recombination leads to a change of 244 TFBSs (Supplementary Table 6). These results suggested the recombination events may cause the change of binding of hundreds of transcription factors at 5’ LTR.

Then, the transcriptional factors bound at 5’ LTR before and after recombination was further analyzed by comparison of the parent strain with the progeny recombinant. Specially, we found that although there were 156 common proteins before and after recombination between the CRF07_BC and the subtype B (the recombinant HB020068, HB070040, and HB070063), these proteins showed different binding position within 5’ LTR after the recombination (Fig. 7a, Supplementary Table 4). Regarding to the other 3 recombinants, HB010151 and HB010161 was separately compared with the parent strain HB010165. As expected, the transcriptional factors bound at 5’LTR was quite similar between the two recombinant, HB010151 and HB010161. However, the association of 58 transcriptional factors with 5’ LTR were completely diminished after the recombination in both HB0151 and HB010161 (Fig. 7b, Supplementary Table 5), in comparison to the parent strain HB010165. Besides, there were only 25 common proteins associated with 5’LTR between the recombinant HB030133 and its parent strain HB070068. However, the location of these 25 proteins within 5’LTR were also changed after the recombination (Fig. 7c, Supplementary Table 6). Among these binding proteins at 5’LTR after recombination, some are very important transcription factors such as AP-1, p53, SP1, P300, and NF-κB (Table 4-6). Together, these data further demonstrated the usefulness of this newly constructed 5’LTR classification system, and highlighted the importance of 5’ LTR characterization in the study of HIV-1 classification, evolution, and biological characterization.

**Figure 7.**
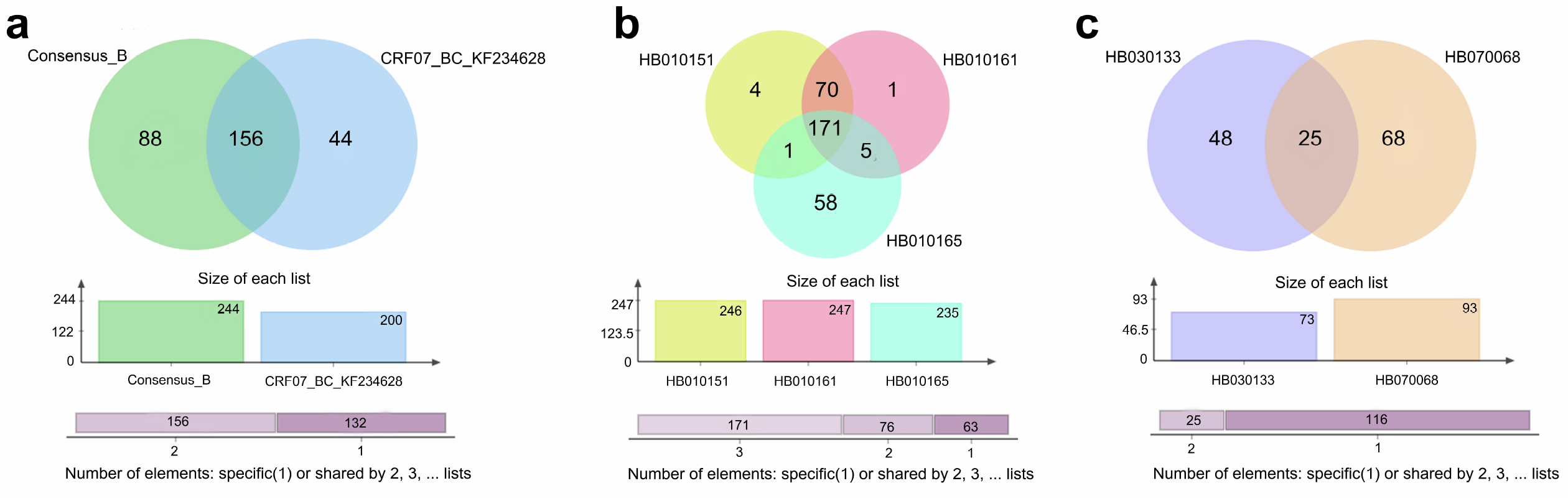
Venn diagram representation of the significant changes of transcription factors caused by recombination at 5’LTR. (a) Change of transcription factors via recombination between the CRF07_BC and the subtype B at 5’ LTR (the recombinant HB020068, HB070040, and HB070063). The green indicates specific transcription factors of subtype B. The blue indicates specific transcription factors of CRF07_BC. The overlap indicates transcription factors shared by subtype B and CRF07_BC. (b) Change of transcription factors via recombination between the recombinant HB010151, HB010161, and the parent strain HB010165 at 5’ LTR. The yellow indicates specific transcription factors of HB010151. The pink indicates specific transcription factors of HB010161. The cyan indicates specific transcription factors of HB010165. The overlap of yellow and pink indicates transcription factors shared by HB010151 and HB010161. The overlap of pink and cyan indicates transcription factors shared by HB010161 and HB010165. The overlap of yellow and cyan indicates transcription factors shared by HB010151 and HB010165. The overlap of yellow, pink, and cyan indicates transcription factors shared by HB010151, HB010161, and HB010165. (c) Change of transcription factors via recombination between the recombinant HB030133 and its parent strain HB070068 at 5’ LTR. The purple indicates specific transcription factors of HB030133. The wheaten indicates specific transcription factors of HB070068. The overlap indicates transcription factors shared by HB030133 and HB070068.

## Discussion

The classification of a virus is rather critical, having a significant impact on molecular epidemiology, basic mechanism, and clinical science. The 5’ LTR accounts for only 6.52% of the full-length HIV-1 genomic RNA but plays crucial roles in the whole life cycle, including the reverse transcription, integration, and transcriptional regulation of HIV-1. In particular, the HIV-1 5’ LTR possesses multiple promoters and enhancers that can be used to drive virtual transcription, and the biological differences of 5’ LTR of some different strains have been described (18, 26). However, the research on 5’ LTR lags far behind that on internal genes, such as *gag, pol*, and *env*, which was restricted by the fact that a systematic classification system of HIV-1 5’ LTR has never been established, although some studies have performed indirect LTR characterization by using the internal gene classification (2, 17-19, 26). This may underestimate the effects of LTR diversity on HIV-1 characterization and evolution.

Thus, this study has proposed a unified and comprehensive classification of HIV-1 5’ LTRs and explored the effect of classification-based 5’ LTR genetic variation on the evolution and biological characteristics of HIV-1. To ensure the reference construction, the 4 recognized formal rules for the subtype nomenclature system were strictly followed. By applying the classification system to 22 Chinese epidemic strains, the classification has been proven practical applicability, revealing some interesting results unclear previously. Specifically, we found the occurrence of large-scale recombination within 5’ LTR of these strains, which has caused the binding change of many important transcriptional factors and most likely would affect the transcriptional activity of 5’ LTR.

The remarkable genetic variation and population plasticity is a prominent feature of HIV-1 (60). This genetic variation originates from two sources, including point mutations and genetic recombination (6, 51). It has been found that frequent genetic recombination existed within HIV-1 genome (6, 37, 38). Our previous work updated the molecular model on HIV-1 genetic recombination (51), which has proposed that a large number of micro-recombination were considered as mutations because they cannot be detected due to the limitation of detection tool strategies and capabilities (51). Compared with mutation, recombination has a much greater remodeling effect on genome structure, as it can lead to a large fragment exchange. For decades, recombinant analysis has revealed 134 CRFs and countless URFs. However, few studies have focused on the recombination analysis within 5’ LTR regions because of the lack of systematic reference sequences. This study has demonstrated that our classification system effectively improves the characterization of HIV-1 5’ LTRs.

In recent years, CRF01_AE and CRF07_BC have been found as dominant HIV-1 strains in China. The discovery of new recombinants within the 5’ LTR region suggested that the prevalence of HIV-1 strains in China could result in the recombination of CRF01_AE and CRF07_BC subtypes. This may generate new recombinants carrying the parental advantages and developing into new epidemic strains. Meanwhile, it also reminded us that the transmission of HIV-1 in China tends to be more complex, while the prevention, control, and treatment of HIV-1 are facing great challenges.

A limitation of the present work is that there are still no classified references for all groups, subtypes, sub-subtypes, and CRFs. With the increase of HIV-1 5’ LTRs and complete sequences obtained, the reference classification system will be improved as the great enrichment of sequences in the HIV sequence database. As the increasing investigation on 5’ LTRs diversity and characterization, we will get a deeper understanding of HIV-1 transmission, evolution, and the basic mechanism of transcriptional regulation. Therefore, our study has proposed a comprehensive classification for HIV-1 5’ LTR, which will facilitate the research on molecular epidemiology, virus detection, and diagnosis, clinical treatment, drug and therapy development.

## Acknowledgments

This study was supported by grants from National Natural Science Foundation of China (31900157, 31900132), the State Key Laboratory of Pathogen and Biosecurity (AMMS).

## Author Contributions

Research design: L.L and L.J. Performed the analysis: X.G, D.Y, L.J, M.L. Sequence collection: H.L, M.C, X.Z, X.W, B.Z, Y.W, C.Y, C.W, Y.L, J.H, X.W, T.L, and J.L. Contributed to the composition of the manuscript: X.G, D.Y, L.J, and L.L.

