## Supplementary Table 1 for "A unified classification system for HIV-1 5’ long terminal repeats"

| Supplementary Table 1 Primers used for amplification of the 5' R-U5-NCR and 3' U3-R regions of the HIV-1 genome | | | | | |
| --- | --- | --- | --- | --- | --- |
| Regions | Primers | Reaction | Sequence (5' to 3' ) | Position^#^ | Length (bp) |
| 3' U3-R | 3R-OF | 1st PCR | CGCAGRTACCTYTAAGACCAATGAC | 9012-9035 | 604 |
|  | 3R-OR |  | ATTGAGGCTTTAAGCAGTGGGTT | 9593-9614 |  |
|  | 3R-IF | 2nd PCR | GGACTGGAWGGGYTAATTTACTC | 9082-9104 | 504 |
|  | 3R-IR |  | YAGCCAGAGAGCTCCCRGGCTC | 9564-9585 |  |
| 5' R-U5-NCR | 5R-OF | 1st PCR | GGTCTCTCTWGKTAGACCAGRTCTGAGCC | 454-483 | 417 |
|  | 5R-OR |  | TTTCTTTCCCCCTGGCCTTAACC | 848-870 |  |
|  | 5R-IF | 2nd PCR | GAGCCYGGGAGCTCTCTGGCTR | 479-500 | 382 |
|  | 5R-IR |  | CCCTGGCCTTAACCNAATTTTNTCCCAT | 834-861 |  |

^#^Position according to the K03455 coordinates
